# Developmental instability of CA1 pyramidal cells in Dravet Syndrome

**DOI:** 10.1101/2022.09.12.507264

**Authors:** Steffan P. Jones, Nathanael O’Neill, Sharon Muggeo, Gaia Colasante, Dimitri M. Kullmann, Gabriele Lignani

**Affiliations:** Department of Clinical and Experimental Epilepsy, UCL Institute of Neurology, University College London, London, UK; San Raffaele Scientific Institute, Via Olgettina 58, 20132, Milan, Italy

**Author notes:** Correspondence to: Gabriele Lignani.

## Abstract

Dravet Syndrome (DS) is mostly caused by heterozygous loss-of-function mutations in the voltage-gated sodium channel *SCN1A* (Na_v_1.1), thought to result in severe epilepsy and neurodevelopmental impairment due to reduced interneuron excitability. Recent studies in mouse models suggest that an “interneuronopathy” alone does not completely explain all the cellular and network impairments seen in DS. Here, we investigated the development of the intrinsic, synaptic, and network properties of CA1 pyramidal cells in a DS model prior to the appearance of overt seizures. We report that CA1 pyramidal cell development is disrupted by loss of *Scn1a*, and propose that this is explained by a period of reduced intrinsic excitability in early postnatal life, during which *Scn1a* is normally expressed in hippocampal pyramidal cells. We also use a novel *ex vivo* model of homeostatic plasticity to show an instability in homeostatic response during DS epileptogenesis. This study provides evidence for an important role of *Scn1a* haploinsufficiency in pyramidal cells in the pathophysiology of DS.

## Introduction

Mutations of voltage-gated sodium channels (VGSCs) are a major cause of neurological disease (Meisler et al., 2021), with both gain- and loss-of-function generating a range of neurodevelopmental disease phenotypes that include autism spectrum disorder (Ben-Shalom et al., 2017; Spratt et al., 2019; Weiss et al., 2003), intellectual disability (Begemann et al., 2019; Liu et al., 2019; Wagnon et al., 2017), and epilepsy (Claes et al., 2001; Escayg et al., 2000; Estacion et al., 2014; Ohmori et al., 2006; Reynolds et al., 2020). One of the commonest diseases caused by mutations in VGSCs is Dravet Syndrome (DS), a severe developmental epileptic encephalopathy, typically associated with *de novo* heterozygous loss-of-function mutations in *SCN1A* (encoding the α-subunit of the voltage-gated sodium channel Na_v_1.1) in 70-90% of patients (Marini et al., 2011; Rosander and Hallböök, 2015). DS is characterised by severe drug-resistant seizures (Chiron and Dulac, 2011; Wirrell, 2016) that display an early age of onset relative to other childhood epilepsies (Cetica et al., 2017; Dravet, 2011), along with a range of other behavioural and developmental comorbidities (Jansson et al., 2020; Li et al., 2011; Zuberi et al., 2022).

Mice carrying heterozygous null or missense loss of function *Scn1a* mutations recapitulate much of the core DS phenotype and have shed light on the cellular and circuit mechanisms underlying DS epileptogenesis. These mice display spontaneous and temperature-sensitive seizures starting around postnatal week 3, and, depending on mouse strain background, a mortality rate of 50% by postnatal day 100 (P100) (Miller et al., 2014; Yu et al., 2006). DS mouse models point to GABAergic interneuron dysfunction as the key driver of network hyperexcitability, with GABAergic interneurons displaying reduced sodium current density (Yu et al., 2006) and a hypoexcitable firing phenotype in all cortical and hippocampal interneuron subtypes investigated thus far (Almog et al., 2021; Goff and Goldberg, 2019; Ogiwara et al., 2007; Tai et al., 2014). Crucially, *Scn1a* hemizygosity restricted to interneurons is sufficient to recapitulate the DS phenotype seen in *Scn1a^+/-^* mice (Cheah et al., 2012). Interneuron hypoexcitability emerges gradually during postnatal development, consistent with normal upregulation of *Scn1a*, which starts at around P10 and does not plateau until P30 (Cheah et al., 2013). The developmental profile of *Scn1a* is well supported by longitudinal electrophysiology studies that show that interneuron excitability is unaffected by *Scn1a* hemizygosity until P18-20, immediately prior to the emergence of seizures (Almog et al., 2021; Favero et al., 2018; Yu et al., 2006). Despite the evidence that interneuron dysfunction is essential for DS epileptogenesis, recent work has revealed that overt interneuron hypoexcitability is restricted to a relatively brief window of postnatal development (Favero et al., 2018) and that patterns of spontaneous interneuron activity recorded *in vivo* are actually relatively normal, in contrast to the loss of excitability shown in *ex vivo* whole cell recordings (De Stasi et al., 2016). These findings suggest that the immediate effects of interneuron hypoexcitability alone are insufficient to explain the lifelong seizures experienced by *Scn1a* heterozygous mice. Further investigation is necessary to understand how transient interneuron hypoexcitability alters brain development, in order to generate an epileptic network that persists beyond that transient phase.

The mechanisms underlying epileptogenesis may extend beyond interneuron dysfunction. Disruptions to glutamatergic or GABAergic transmission do not occur in isolation to one another, with changes to one neurotransmitter system capable of generating compensatory changes in the other (He et al., 2018; Howard et al., 2014; Swanwick et al., 2006). This general principle is thought to maintain the fine balance between excitation and inhibition that emerges by the late stages of postnatal development (Adesnik, 2018; Bhatia et al., 2019; Mitchell and Silver, 2003; Zhou and Yu, 2018). Experimental induction of interneuron death using *Dlx*^(-/-)^ mice, for instance, generates homeostatic reductions in both the frequency of miniature excitatory postsynaptic currents (mEPSC) and in the intrinsic excitability of pyramidal cells in the hippocampus, as reflected in a decrease in input resistance and an increase in rheobase resulting in an overall decrease in action potential generation (Howard et al., 2014). Such studies suggest that loss of inhibitory transmission in DS during postnatal development may alter the properties of the pyramidal cells that they innervate, potentially contributing to the emergence of a network capable of generating spontaneous seizures.

Early work suggested that pyramidal cells are not directly affected by *Scn1a* mutations. Thus, *Scn1a* hemizygosity was associated with no change in pyramidal cell sodium current density (Yu et al., 2006). This is consistent with sodium currents in pyramidal cells being primarily mediated by *Scn2a* (Na_V_1.2) and *Scn8a* (Na_V_1.6) (Royeck *et al*., 2008). Around the onset of seizures (P20-25) and at later ages (P33-45), CA1 pyramidal cell excitability is unchanged by loss of *Scn1a* relative to wildtype. However, at an earlier (pre-seizure) developmental stage (P14-16), recent studies have reported a slight increase in maximal firing frequency of CA1 pyramidal cells during current injection in *ex vivo* experiments (Almog et al., 2021; Chancey and Howard, 2022). Changes in intrinsic and synaptic properties of CA1 pyramidal cells have also been reported after the onset of seizures in DS mice (Almog et al., 2022; Dyment et al., 2020), with more substantial changes in glutamatergic neurons noted in other subcortical regions (Studtmann et al., 2022; Yan et al., 2021). Despite the deficits in GABAergic transmission, no alterations in synaptic integration have been reported in CA1 pyramidal neurons in adult DS animals (Chancey et al. 2022).

Given the emerging evidence for subtle alterations in excitatory neuron properties in DS models, we undertook the present study to investigate the synaptic and intrinsic properties of CA1 pyramidal cells at three timepoints across the epileptogenic period (P10-20) in *Scn1a* hemizygous mice, in order to better understand how the hypoexcitability of interneurons that emerges during this period affects the development of the excitation/inhibition balance (E/I balance) in the hippocampus. We asked if GABAergic hypofunction generates compensatory changes to the intrinsic and/or synaptic development of CA1 pyramidal cells. We found that, early (P10-12) stages of epileptogenesis are characterized by pyramidal cell hypoexcitability, seemingly due to an early phase of postnatal life in which *Scn1a* expression is comparatively elevated in pyramidal cells. We also uncovered an E/I imbalance in CA1 pyramidal neurons in response to Schaffer collateral stimulation by P18-20, due to impaired feedforward inhibition (FFI). Furthermore, several changes to both intrinsic and synaptic properties emerge at P18-20, consistent with a homeostatic compensation for impaired inhibition. Finally, homeostatic compensations for increased network activity were severely disrupted at P10-12 and P14-16, and stabilised by P18-20, when comparing DS to WT mouse hippocampus. These results reveal extensive and dynamic pathological processes accompanying the emergence of the DS phenotype, involving developmental changes to the intrinsic properties of excitatory neurons.

## Results

### Hippocampal E/I imbalance manifests by postnatal day 18 in Dravet syndrome mice

To understand how interneuron hypofunction relates to disinhibition of CA1 pyramidal cells we assessed the E/I balance in *ex vivo* brain slices (Zhou and Yu, 2018) across the epileptogenic window in *Scn1a^+/-^* (DS) mice and their control (WT) littermates (**Fig.1; Fig. S1**). We recorded Schaffer collateral-evoked monosynaptic excitatory postsynaptic currents (evoked EPSCs, eEPSCs) and disynaptic inhibitory postsynaptic currents (evoked IPSCs, eIPSCs) in CA1 pyramidal cells at three ages: P10-12, P14-16 and P18-20 (**Fig.1A,B**).The E/I ratio (eEPSC/eIPSC amplitude ratio) showed divergent trends between genotypes: the WT E/I ratio decreased progressively with development; in contrast, the DS E/I ratio shifted progressively in favour of excitation. By P18-20, the E/I ratio was significantly greater in DS than WT neurons (**Fig.1C**). Both eEPSC and eIPSC amplitude showed a progressive increase between early (P10-12) and late (P18-20) phases of epileptogenesis, in both WT and DS slices. Despite increasing in both genotypes, eIPSC amplitudes displayed a significant genotype-dependent decrease in DS compared to WT cells (**Fig.1D**). We also assessed the probability of successfully evoking eEPSCs and eIPSCs. The eIPSC success probability displayed a significant reduction across time in DS cells, whilst that of eEPSCs was unaffected (**Fig.1E**).

**Figure 1.**
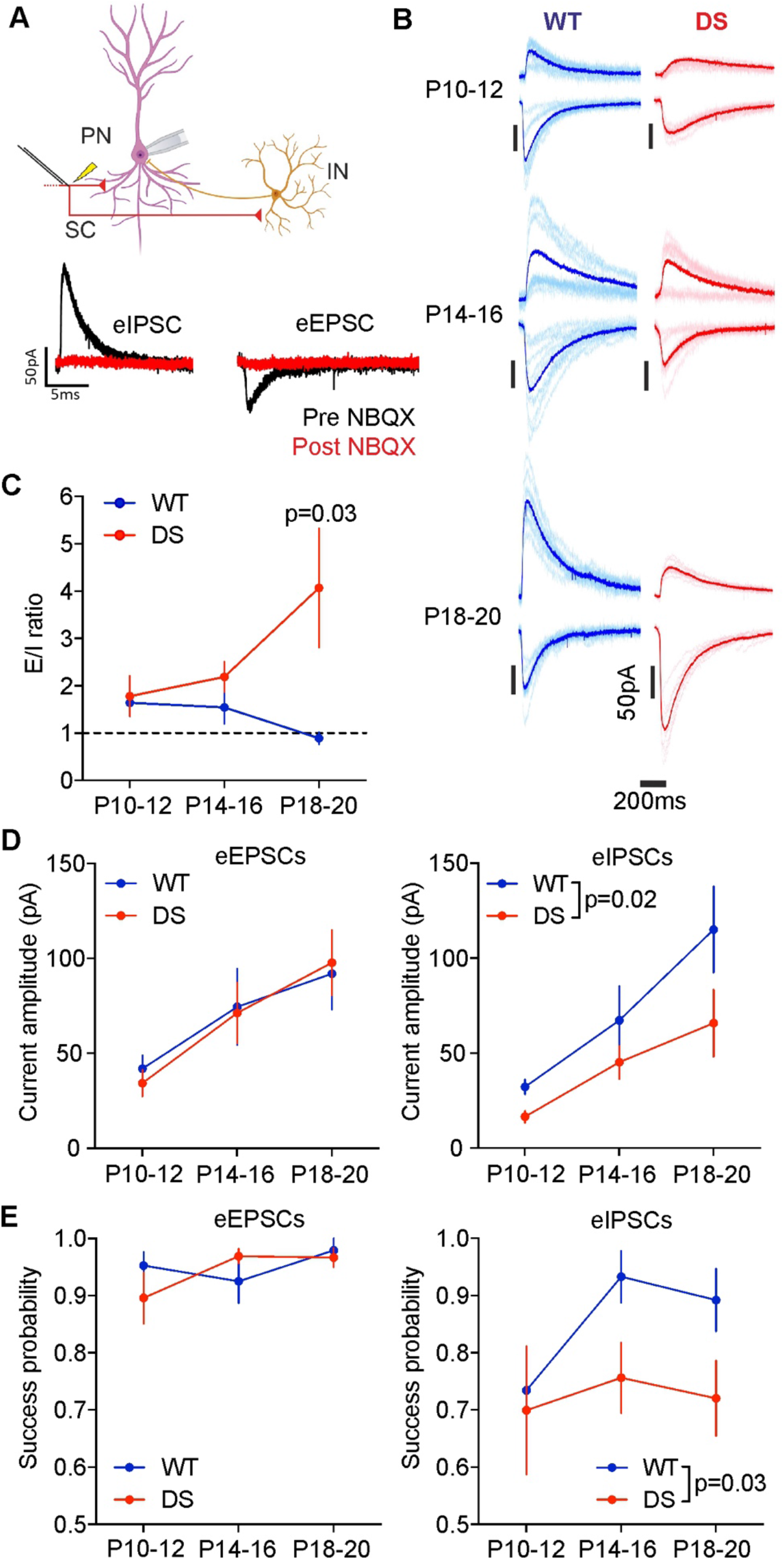
Interneuron hypofunction in DS results in hippocampal E/I imbalance at the Schaffer collateral synapses by P18-20. **A.** Experimental design for evoked current recordings in CA1 pyramidal neurons. Both monosynaptic EPSCs and disynaptic IPSCs were evoked via electrical stimulation of stratum radiatum to evoke Schaffer collateral mediated currents. NBQX application prevents both monosynaptic eEPSCs and disynaptic eIPSCs. **B.** Representative traces of both eEPSCs and eIPSCs in WT (blue) and DS (red) mice at P10-12, P14-16 and P18-20. **C.** E/I ratio across development. Two-way ANOVA followed by Holm-Sidak’s multiple comparisons test. **D.** eEPSC and eIPSC current amplitudes across development. Two-way ANOVA. **E.** Probability of evoking eEPSCs and eIPSCs across development. Two-way ANOVA. n = WT [31, 16, 12]; DS [10, 21, 30] cells, N = WT [8, 6, 6]; DS [4, 5, 8] mice.

### Developmental changes in spontaneous synaptic activity are perturbed during DS epileptogenesis

Given the disruption to evoked inhibitory transmission, we asked whether excitatory and/or inhibitory spontaneous activity was affected during development in DS mice. We quantified spontaneous excitatory and inhibitory postsynaptic currents (sEPSCs and sIPSCs, respectively), from traces collected in between evoked synaptic currents (**Fig.2A**). sEPSC activity, measured as the integral of sEPSCs, was initially comparable between genotypes at P10-12, before becoming significantly elevated at P14-16 in DS cells relative to WT (**Fig.2B**). Whilst excitatory synaptic activity continued to increase between P14-16 and P18-20 in WT mice, in DS mice it decreased between P14-16 and P18-20, such that it was significantly lower in DS mice compared to WT at P18-20 (**Fig.2B**). The overall trajectory of spontaneous excitatory activity demonstrates a period of transient hyperactivity that is subsequently downregulated just prior to the anticipated age of seizure onset, coincident with the emergence of temporal E/I imbalance due to inhibitory hypofunction (**Fig.1C**). The developmental pattern of sIPSC charge transfer followed a similar pattern to the emergence of evoked IPSC amplitude, with DS activity nearly identical to WT at the two younger age groups, before a 20.5% reduction in sIPSC charge transfer in DS cells relative to WT cells at P18-20 (**Fig.2B**).

**Figure 2.**
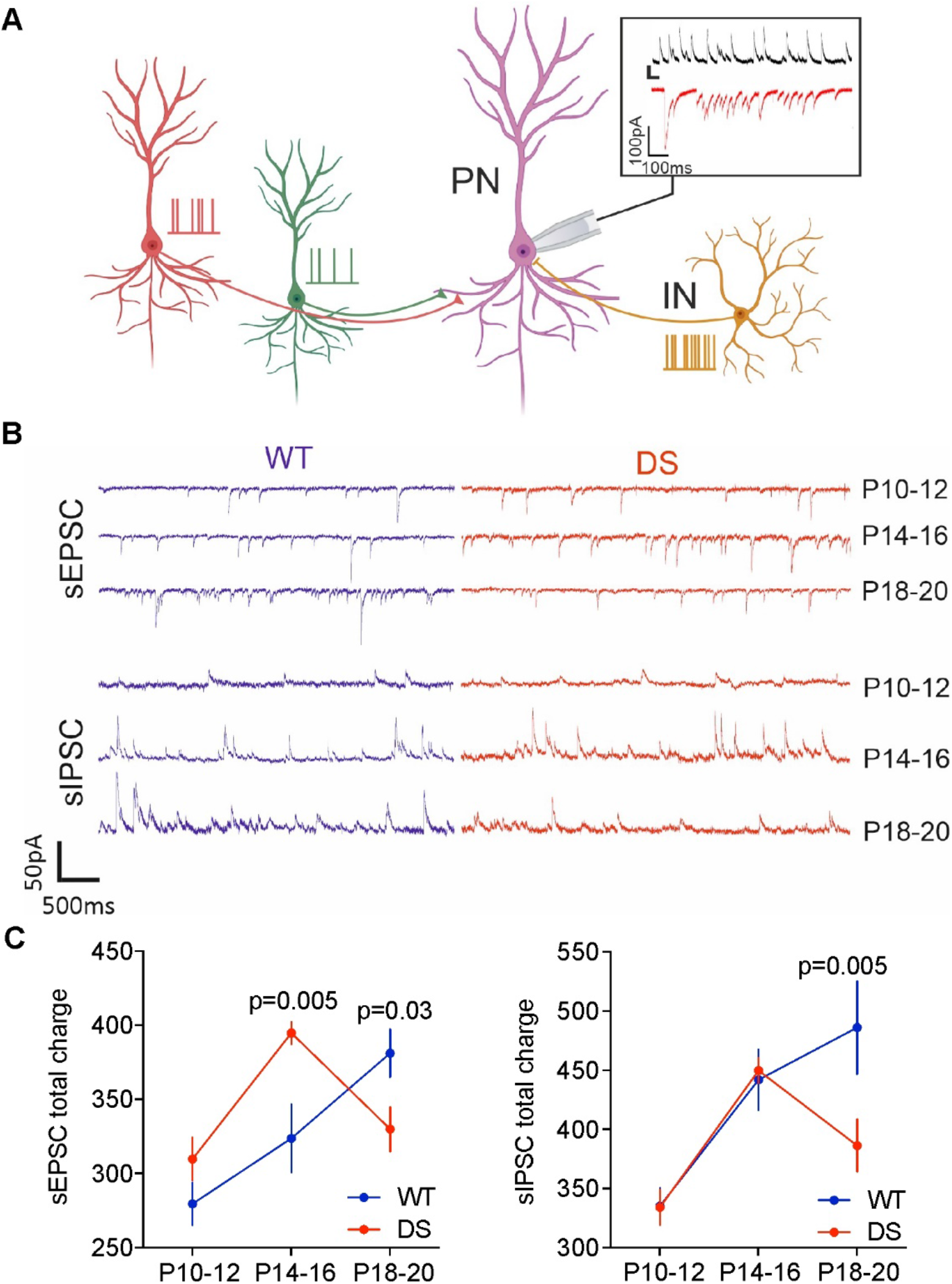
Spontaneous EPSC and IPSC charge transfer to CA1 pyramidal neurons are altered across the time course of DS epileptogenesis. **A.** Spontaneous firing of both pyramidal cells and interneurons generates spontaneous synaptic activity *ex vivo*. Spontaneous currents that occurred between slice stimulation were quantified by total charge transfer. **B**. Representative periods of spontaneous activity across all age groups for both WT and DS. **C.** Total sEPSC and sIPSC charge transfer in WT and DS mice at P10-12, P14-16 and P18-20. Two-way ANOVA followed by Holm-Sidak’s multiple comparisons test. n = WT [51, 33, 36]; DS [23, 46, 57] cells, N = WT [8, 6, 6]; DS [4, 5, 8] mice

### DS CA1 pyramidal cells display an unexpected reduction in intrinsic excitability during early epileptogenesis

The emergence of interneuron hypoexcitability during the epileptogenic period is well characterized in DS mice (Cheah et al., 2013; Favero et al., 2018), but the direct or indirect effects of *Scn1a* heterozygosity on pyramidal cell excitability remain unclear. We carefully analysed intrinsic excitability properties across the same age groups during epileptogenesis (**Fig.3A,B**). At P10-12 we observed an unexpected significant reduction in pyramidal cell excitability as quantified by current step injection inputoutput curve and maximal firing frequency (**Fig.3C,F**). Despite the E/I imbalance progressively shifting in favour of excitation from P14-16 onward in DS mice, the hypothesized compensatory alterations to pyramidal cell excitability were not seen at either P14-16 or P18-20. No significant differences in input output curve were seen, although a slight increase in maximal frequency occurred at P14-16, as previously reported (Almog et al., 2021). No significant difference was noted in overall maximal frequency at P18-20. No differences between genotypes were observed in either input resistance or cell capacitance in any age group (**Fig.S2**). Overall excitability data suggests that the epileptogenic phase of DS mice is characterized by an unanticipated instability in the intrinsic properties of CA1 pyramidal cells, most notably during the earliest phases of DS epileptogenesis in which no interneuron dysfunction is known to occur.

**Figure 3.**
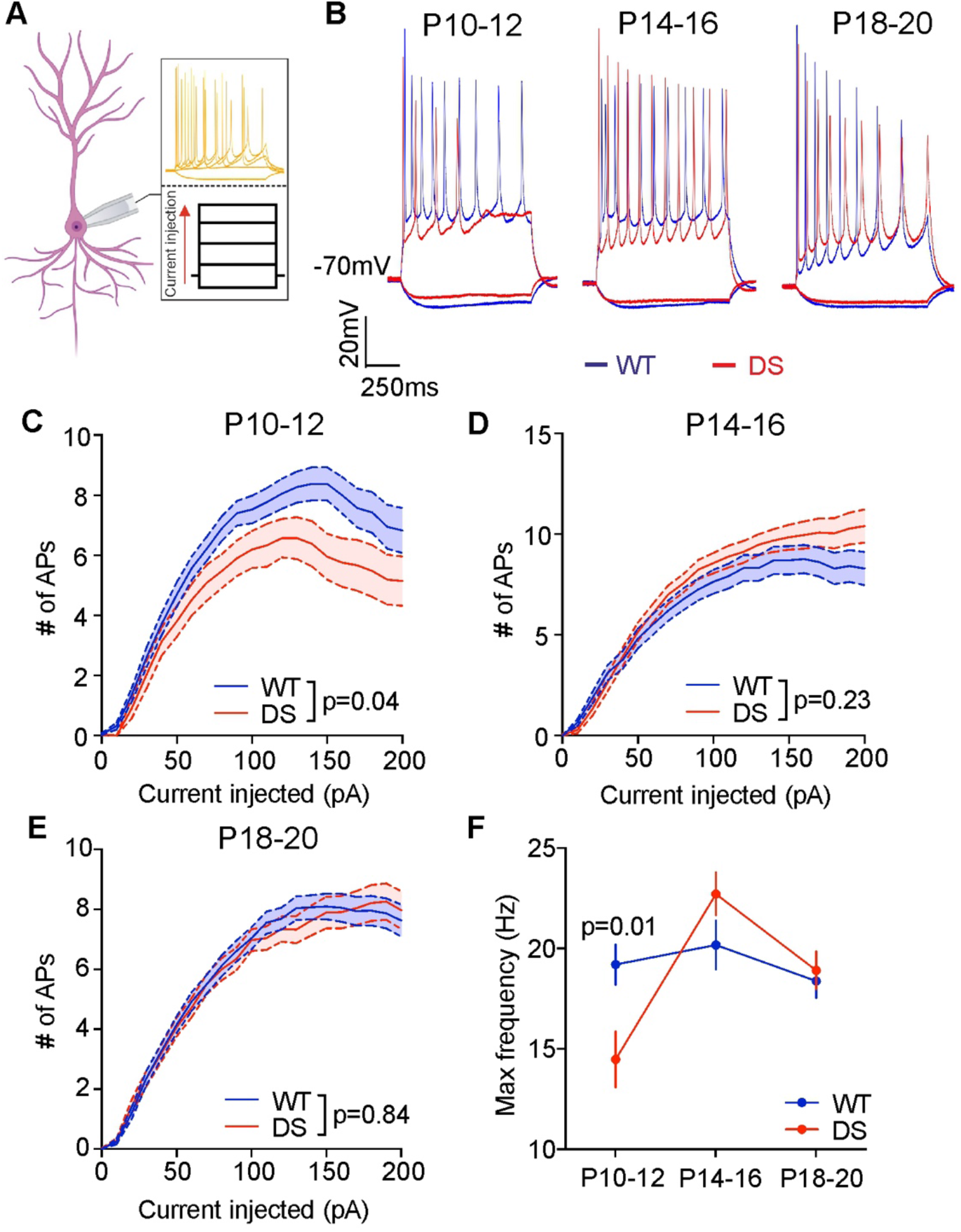
CA1 pyramidal cells exhibit unanticipated hypoexcitability at P10-12 in DS mice. **A.** Current injection-evoked action potential firing was recorded from CA1 pyramidal cells. Square pulse current steps were injected between −40 and +200pA in 10pA increments. **B.** Representative 100pA square-pulse evoked action potentials for WT (blue) and DS (red) at P10-12, P14-16 and P18-20. **C-E.** Input-output curves for WT and DS cells at P10-12 (**C**), P14-16 (**D**) and P18-20 (**E**). **F.** Maximal frequency reached for WT and DS at P10-12, P14-16 and P18-20. All graphs: two-way ANOVA followed by Holm-Sidak’s multiple comparison test. n = WT [40, 34, 52]; DS [21, 25, 46] cells, N = WT [7, 5, 8]; DS [6, 7, 7] mice.

### Scn1a is expressed in CA1 pyramidal cells in early development

We asked whether the decrease in excitability of CA1 pyramidal cells can be ascribed to the expression of *Scn1a* in hippocampal excitatory neurons at P10-12. We dissected hippocampi of GAD67-GFP mice at P0 and P10 and isolated excitatory neurons (**Fig.4A.B**). Using qPCR we observed clear expression of *Scn1a* in excitatory neurons at both ages (**Fig.4C**). Interestingly, *Scn1a* expression was highest at P0, as previously reported (Liang et al., 2021), but was also elevated at P10 (**Fig.4C**). Unfortunately, technical challenges in this protocol produced unreliable data for excitatory neurons at P20. However, *Scn1a* expression in total neurons at P20 was higher than at P10, in line with the increased *Scn1a* expression in inhibitory neurons (Cheah et al., 2013).

**Fig.4.**
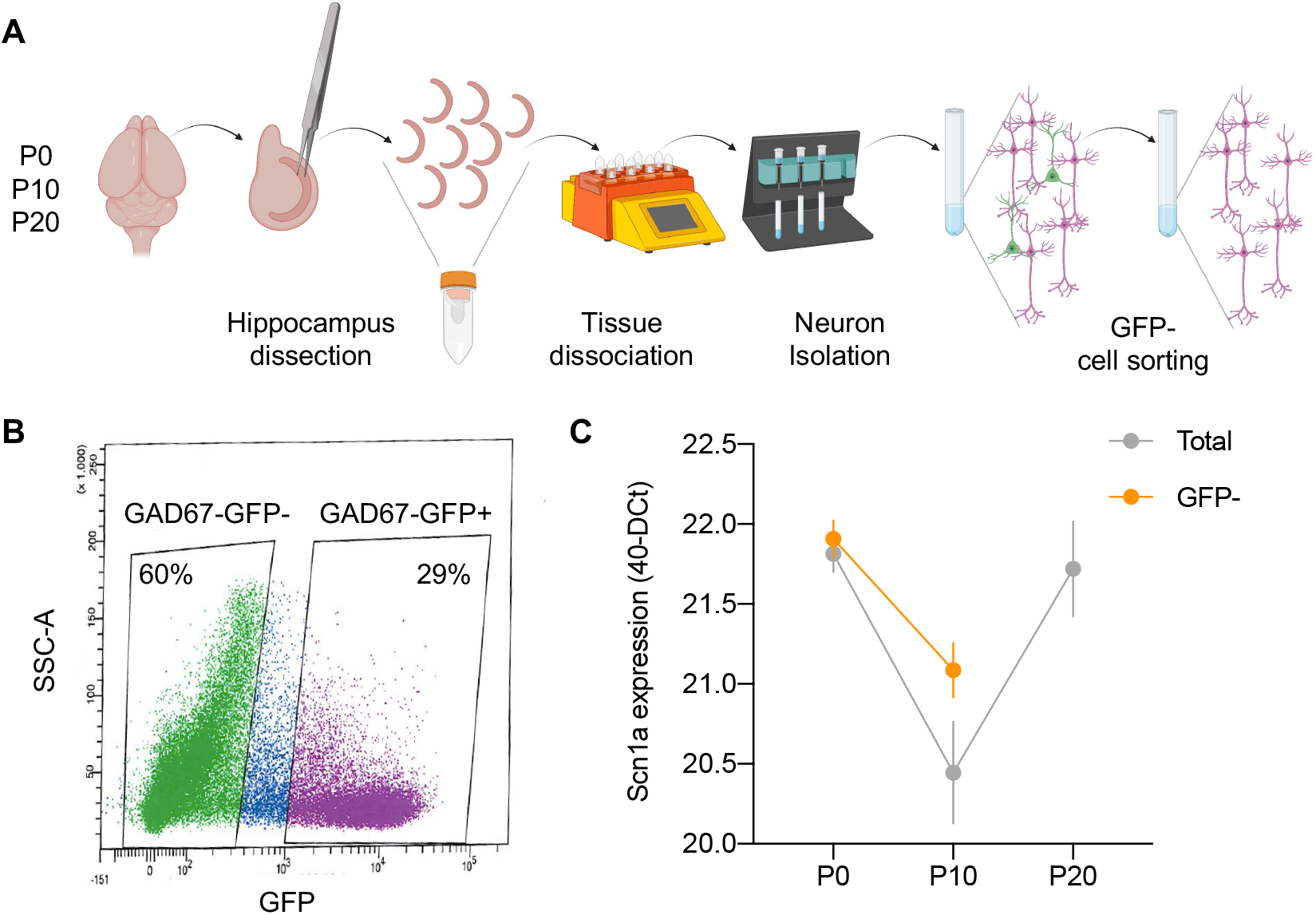
*Scn1a* is expressed in excitatory neurons early in development. **A.** Experimental procedure for separation of GAD67-GFP negative neurons from the hippocampus. **B.** Representative scatter plot for the FACS sorting of GAD67-GFP negative neurons. **C.** Scn1a expression in either Total neuronal (grey) or GAD67-GFP negative (orange) fraction expressed as 40 (cycles) – DCt. Total neurons: n=[2,2,2] reactions; N=[2,2,6] pulled animals for each reaction. GAD67-GFP negative: n=[2,2] reactions; N=[4,4] pulled animals for each reaction.

### Pyramidal cell action potential parameters are unchanged by Scn1a heterozygosity

Single action potential (AP) properties are indicative of changes in membrane ion channel composition and can therefore be used to indicate underlying mechanisms of altered excitability seen across epileptogenesis in DS mice. We applied current slope injections to CA1 pyramidal cells to assess these properties in rheobase-evoked action potentials (**Fig. S3**). Although almost all parameters assessed exhibited age-related changes, these changes were largely identical between genotypes (**Fig. S3**).

### Sag potentials are enhanced in DS CA1 pyramidal cells at P18-20

We also analysed hyperpolarizing current-evoked sag potentials, indicative of the activation of hyperpolarization-activated cyclic nucleotide–gated (HCN) channels (Biel et al., 2009) (**Fig.5A**). At each age group, the amplitude of sag potentials showed a linear relationship with the amplitude of hyperpolarizing current injection and this period of postnatal development saw a gradual decrease in sag potential amplitude in both genotypes between each age group (**Fig.5B-D**). No significant differences between sag potentials or associated linear fits were seen at either P10-12 or P14-16 (**Fig5B,C**). However, at P18-20, we observed a significant difference between both raw data and fitted slopes (**Fig.5D**). This data suggests that HCN channel function and/or expression levels remain comparatively elevated in DS mice compared to their WT littermates by P18-20.

**Figure 5.**
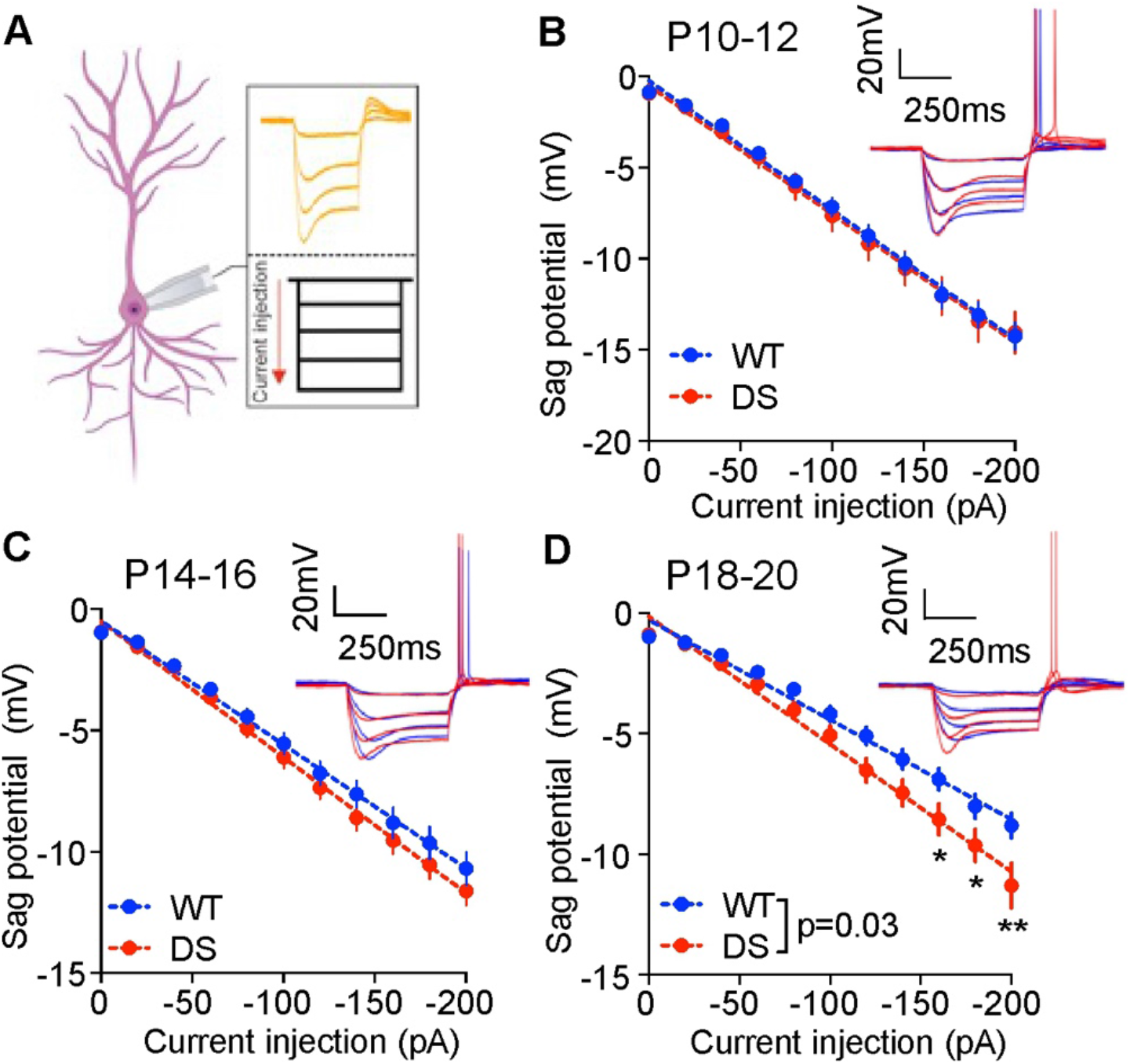
Sag potential amplitudes are elevated in DS CA1 pyramidal cells at P18-20. **A.** Sag potentials were evoked through 500ms square pulse hyperpolarizing current steps from 0 to −200pA increasing in 20pA increments. **B – D.** Sag potential amplitudes in WT and DS mice at both P10-12 **(B)**, P14-16 **(C)** and P18-20 (**D)**. All datasets are fitted with a linear slope. Two-way ANOVA followed by Holm-Sidak’s multiple comparison. *p<0.05; **p<0.01. n = WT [40, 34, 50); DS [21, 25, 46] cells, N = WT [7, 5, 8]; DS [6, 7, 7] mice.

### Input integration is unaltered by E/I imbalance during DS epileptogenesis

The integration of different incoming synaptic inputs is crucial to neuronal activity, as APs are typically generated via the summation of multiple inputs rather than by activity at a single synaptic connection. One of the key factors in determining the temporal window of input summation is disynaptic inhibition (Magee, 2000; Poirazi et al., 2003; Pouille and Scanziani, 2001). Having demonstrated that late-stage DS epileptogenesis is characterized by deficits in disynaptic inhibition, we set out to determine how input integration changes across epileptogenesis. To overcome issues introduced by the limited tissue volume available for concurrent electrode placement in juvenile mouse hippocampal slices, we used a combination of Schaffer collateral stimulation and injection of an artificial AMPA-receptor like conductance via dynamic clamp to assess input integration across the epileptogenesis window (**Fig.6A**). We observed no significant differences at any age group comparing WT and DS, either when raw data were analysed or when data were quantified as either area under the curve or by taking the standard deviation (σ) of fitted Gaussian curves, with greater σ values indicating a wider integration window (O’Neill and Sylantyev, 2018) (**Fig. 6B-E; Fig. S4**). However all 3 metrics tend to indicate a widening of AP integration window at both P14-16 and P18-20 (**Fig. 6D,E; Fig. S4)**. Only the direct comparison of fitted curves at P14-16 reveal significant differences between the two genotypes, with fitted curves showing no significant differences at the other timepoints. Fitted curves are shared between genotypes at P10-12 and P18-20 and plotted separately at P14-16. Given the strong evidence that feedforward inhibition is a key regulator of input integration (Pouille and Scanziani, 2001), the E/I imbalance noted at P18-20 (**Fig.1C**) could have been compensated by the decrease in sag potential (**Fig.5D**). Stimulation intensity used during experiments, calibrated via the minimal stimulation intensity needed to consistently evoke APs during calibrated conductance injection, underwent a slight increase in both genotypes during postnatal development but we didn’t observe any genotype-dependent differences (**Fig.S4**). In contrast, conductance injection required to evoke APs consistently decreased with age in WT mice but increased between P14-16 and P18-20 in DS mice (**Fig.S4**).

**Figure 6.**
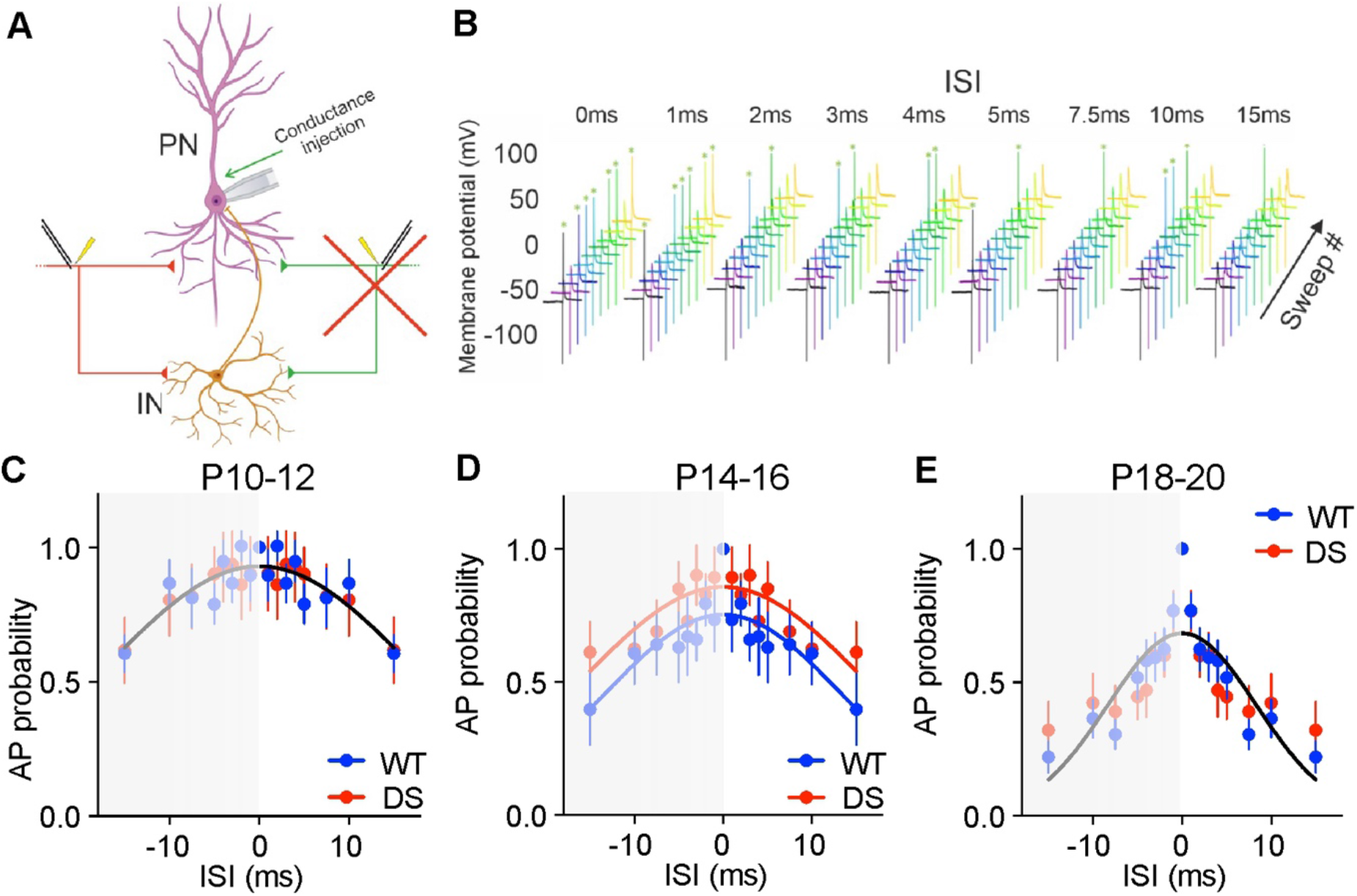
DS mice show stable input integration during development. **A.** The temporal integration window was probed by a mixed protocol that used both stimulation-evoked synaptic input and an artificial dynamic clamp-evoked EPSC-like conductance delivered at an increasing interstimulus interval, from 0 to +15ms. **B.** Representative traces demonstrating the relationship between interstimulus interval (ISI) and action potential probability. **C-E.** Integration properties, as assessed by the relationship between ISI and action potential probability, for WT and DS mice at P10-12 (**C**), P14-16 (**D**) and P18-20 (**E**).Two-Way ANOVA. Each dataset is mirrored (mirrored data behind grey box) to allow a Gaussian curve to be fitted. When fitted curves are directly compared, DS exhibit a widened integration window at P14-16 (**D**, p = 0.003, extra sum of squares F-test). n = WT [11, 8, 19]; DS [10, 16, 11] cells, N = WT [6, 3, 8]; DS [4, 5, 5] mice.

### Homeostatic plasticity is altered in DS mice during development

How perturbations to synaptic or intrinsic physiology interact with homeostatic plasticity is unclear, with recent work attempting to address this question in a variety of neurodevelopmental conditions (Booker et al., 2020; Ellingford et al., 2021). We therefore asked whether the instability to the development of intrinsic and synaptic properties in DS pyramidal cells results in instability of homeostatic response during DS epileptogenesis. We adapted a protocol using pharmacological manipulation of activity in acute *ex vivo* slices (Booker et al., 2020; Ellingford et al., 2021). Here we applied the voltage-gated potassium channel blocker 4-aminopyridine (4AP), a widely used chemoconvulsant that has previously been employed to test the homeostatic responses of cultured cells to elevated activity levels (Pozzi et al., 2013), in order to elevate synaptic and neuronal activity for a defined duration. Firstly, we demonstrated that the increased network activity was sustained throughout the protocol by recording spontaneous APs in CA1 pyramidal cells 3-5 hours after 4AP (all recordings in 4AP) (**Fig.7 A-C**). Then we assessed compensatory changes in intrinsic excitability of CA1 pyramidal cells in slices 6 hours after 4AP application and in matched control slices bathed in standard aCSF across the same time period. All recordings were performed in aCSF after washout of 4AP. All data are presented as 4AP-exposed cell data normalized by subtraction from the mean values of the matched aCSF-exposed control group. We observed a significant difference in post-4AP excitability, with inter-genotype differences becoming more prominent as the injected current amplitude increased, at both P10-12 and P14-16 (**Fig.7D-E**). Interestingly although differences were observed at both ages the sign of the difference was entirely reversed between genotypes, with 4AP-exposed DS cells compensating for the increase network activity at P10-12 and not compensating anymore at P14-16 (**Fig.7D,E**). We observed no significant changes between genotypes at P18-20 (**Fig.7 F**), although both aCSF- and 4AP-treated cells were significantly less excitable in DS slices than in WT (**Fig. S6**). In addition to overall excitability, AP morphology was assessed alongside other parameters relevant to neuronal excitability (**Fig. S7**). Holding current and maximal decay velocity both exhibited genotype-dependent changes. At all 3 age groups, 4AP application resulted in an increase in holding current required to maintain the cell membrane voltage at −70mV in DS slices and a decrease in WT slices, with a significant inter-genotype difference seen at both P10-12 and P14-16 (**Fig. S7**). In line with the overall changes seen in excitability at P10-12, significant divergent changes in both AP initiation threshold and maximal rate of AP decay were seen between genotypes. Although other parameters also correlated with the overall changes seen at P10-12 i.e. maximal rate of AP rise and AP half-width, these changes were not significantly different between genotypes (**Fig. S7**).

**Figure 7.**
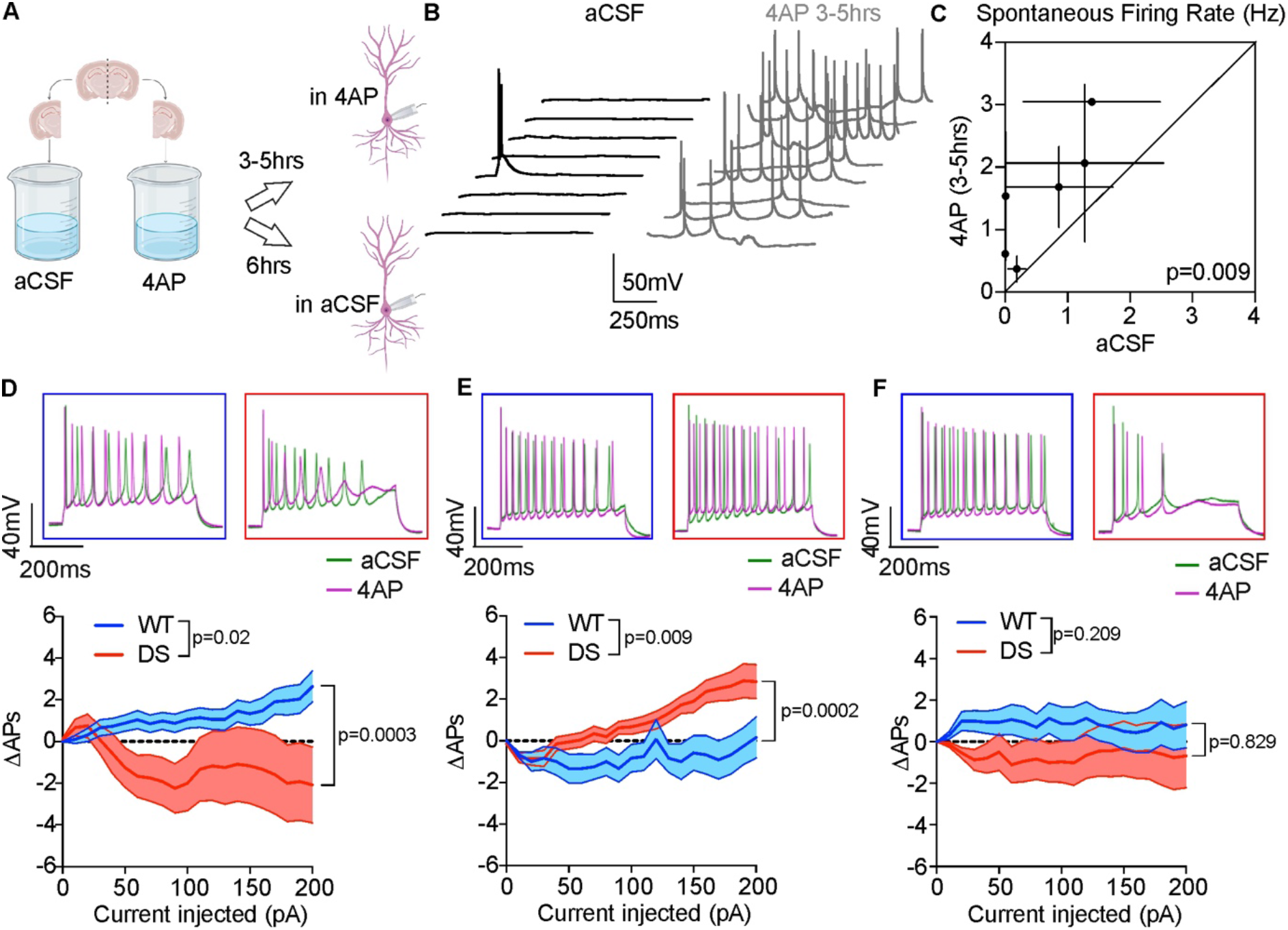
DS pyramidal neurons undergo altered intrinsic homeostatic plasticity in response to prolonged 4AP application. **A.** Slices were bisected and each half exposed to standard aCSF or aCSF + 25μM 4AP for 6 hours. Spontaneous APs, were assessed in WT cells in aCSF and 3-5hrs after 4AP (recorded in 4AP). Intrinsic properties of CA1 pyramidal cells were assessed in aCSF and 6hrs after 4AP (recorded in aCSF). **B**. Representative trace for WT cells in aCSF or 3-5hrs after 4AP. **C.** Plot comparing spontaneous APs in aCSF and 3-5hrs after 4AP in the same animals. N= 6 animals. Paired Student’s T test. **D-F.** Representative trace (*upper*) and difference in number of induced APs (*bottom*) of WT and DS cells in aCSF vs 6hrs 4AP at P10-12 (**D**), P14-16 (**E**) and P18-20 (**F**). Two-Way ANOVA followed by Holm-Sidak’s multiple comparison test. P value of the multiple comparison is the lowest value for the comparison. n = WT [32, 12, 16]; DS [12, 21, 14] cells, N = WT [7, 3, 4]; DS [3, 5, 4] mice.

## Discussion

The main pattern that emerged from this data is that the intrinsic properties of CA1 pyramidal cells are highly unstable over the course of DS epileptogenesis. Multiple changes precede the onset of inhibitory hypofunction and E/I imbalance in DS mice, the main cause so far of DS pathophysiology. The fact that changes to pyramidal cell properties precede the onset of interneuron dysfunction suggests that *Scn1a* haploinsufficiency has an early and direct impact on pyramidal cell function in the murine hippocampus during early postnatal life. This is supported by qPCR data which demonstrates expression of *Scn1a* at early postnatal timepoints and brings the role of non-GABAergic cell types in mediating *Scn1a* mutation-associated pathophysiology into greater prominence. The earliest alteration to either intrinsic or synaptic physiology that we identified in this study is pyramidal cell hypoexcitability at P10-12, challenging the notion that DS epileptogenesis is driven solely by an ‘interneuronopathy’ (Catterall, 2018). Whilst the qPCR data do not challenge the general idea that *Scn1a* gene expression is higher in interneurons, it reveals that there is Scn1a expression in hippocampal excitatory cell at P0 and P10 (**Fig.4**). This fits with the pattern of pyramidal cell excitability across development in DS mice. A similar rescue of pyramidal cell excitability at P10 would confirm the role of Scn1a deficit in generating pyramidal cells hypoexcitability. The relevance of pyramidal cell *Scn1a* heterozygosity to the overall DS phenotype is questionable as it has been demonstrated that interneuron-selective knockout of Scn1a is sufficient to generate a DS-like phenotype (Cheah et al., 2012). However, although the functional impact of pyramidal cell selective *Scn1a* hemizygosity has been tested, the intrinsic excitability of pyramidal cells was not evaluated. No overt behavioural changes were seen when an Emx1-Cre line was used to selectively delete Na_v_1.1 from pyramidal cells, but importantly the loss of *Scn1a* in pyramidal cells did augment disease progression when combined with *Scn1a* deletion in interneurons via a dual Emx1-Cre, VGAT-Cre mouse line (Dutton et al., 2013). This raises the possibility that the pyramidal cell phenotype that we see is in fact anti-epileptogenic. Although this is supported by P10-12 CA1 hypoexcitability, the re-normalization of pyramidal cell activity levels by the age of seizure onset at P18-20, a phenotype observed by other studies (Almog et al., 2021), suggests that this protection does not result from diminished pyramidal cell excitability by the age of seizure onset. Regardless of pathophysiological relevance, the fact that an impact was seen at all suggests once again that *Scn1a* is functionally expressed by pyramidal cells at a sufficient level to alter Dravet syndrome disease progression.

E/I imbalance in DS mice emerges incrementally between P10 and P20, a time course that matches the progressive emergence of intrinsic hypoexcitability in interneurons (Cheah et al., 2013; Favero et al., 2018). Whilst changes to inhibitory transmission have previously been demonstrated in a wide variety of DS models and through a variety of methods (Almog et al., 2021; De Stasi et al., 2016; Han et al., 2012; Uchino et al., 2021), this study highlights synaptic changes at multiple points throughout the epileptogenic window, assessed as both temporal and global synaptic E/I balance.

Despite the loss of E/I balance and reduced IPSC probability at P18-20, no changes to input integration were detected (**Fig.6**) and spontaneous excitatory charge transfer was in fact reduced rather than increased at this age (**Fig.2**). Using a theta burst protocol, unaltered input integration has also been demonstrated in DS mice aged between P21 and P38 (Chancey and Howard, 2022), supporting the findings reported here. This lack of change in input integration can help explain why *in vivo* network activity levels are largely unaltered in DS mice (De Stasi et al., 2016) and suggests that disinhibition-generated hyperactivity alone is an overly simplistic explanation for the emergence of a seizures in DS mice. It is also possible that homeostatic changes compensate for the loss of disynaptic inhibition to prevent altered integration. Indeed, we noted an upregulation of the HCN-channel mediated sag potential at P18-20 in DS, a channel type known to limit input integration window in CA1 pyramidal cells (Pavlov et al., 2011) and known to be under activity-dependent homeostatic control (Gasselin et al., 2015).

Surprisingly, changes most consistent with network hyperactivity occur at P14-16 with a 25% increase in spontaneous EPSC charge transfer noted. This change precedes E/I imbalance, although evoked IPSC probability is already reduced at this age, and precedes the majority of changes noted to the intrinsic excitability of interneurons (Almog et al., 2021; Favero et al., 2018). Changes to input integration are also most prominent at this age (**Fig.6**). Although a slight non-significant increase in the intrinsic excitability of CA1 pyramidal cells can be seen in this study, found by others to be significant at this age (Almog et al., 2021), this increase only manifests at high amplitude, non-physiological current injections (**Fig.3**). The subsequent reduction in sEPSC charge transfer at P18-20 may itself by a homeostatic response to network hyperactivity noted at P14-16 (**Fig.2**), with homeostatic synaptic scaling of excitatory synaptic strength a widely reported mechanisms by which networks adapt to altered activity (Diering et al., 2017; El-Hassar et al., 2007; Queenan et al., 2018; Turrigiano et al., 1998). Miniature EPSCs, not sEPSCs, would have to be recorded to understand if reduced excitatory synaptic activity results from homeostatic downscaling of synaptic strength.

In this study we also implemented a novel protocol to assess homeostatic responses to altered circuit activity in *ex vivo* brain tissue. In DS mice, the most consistent 4AP-induced changes are to holding current (**Fig.7**), with a consistent increase in 4AP exposed DS slices relative to DS aCSF slices. Since holding current is in part determined by resting membrane potential, these data suggest that 4AP-application induced a resting membrane hyperpolarization in DS cells in all 3 age groups, raising the possibility that greater changes to overall firing curves may have been seen if experiments were conducted at resting membrane potential rather than in cells voltage clamped to −70mV. The most prominent differences between genotypes can be seen in the P10-12 age group. At this age, maximal firing frequency increases in WT mice whilst decreasing in DS mice. Overall, the changes seen at P10-12 suggest that only DS pyramidal cells homeostatically respond to altered activity levels. The direction of change is reversed at P14-16, with the response to hyperactivity in 4AP-exposed DS slices being to increase in intrinsic excitability. This dysfunctional response occurs at a critical period of DS epileptogenesis, as it is the age at which in the switch from Na_v_1.3 to Na_v_1.1 dominance occurs in interneurons (Beckh et al., 1989; Cheah et al., 2013) and is followed by seizure onset typically just days later. It is possible therefore that this response represents a form of positive feedback-loop, with increases in network activity generating further increases in the excitability of pyramidal cells. The effect of 4AP application is more limited at P18-20, with I-O curves not significantly differing between the two genotypes although maximal frequency showed a non-significant decrease in DS cells and an increase in WT cells.

The role of interneuron dysfunction in Dravet Syndrome is indisputable, with clear evidence that *Scn1a* mutations generate epileptic networks primarily due to reductions in the excitability of interneurons (Cheah et al., 2012). However, both this study and other recent investigations into DS circuits suggest that *Scn1a* mutations do not exclusively affect interneurons (Almog et al., 2021). Overall, these data revealed that DS epileptogenesis does not start at the point at which interneuron *Scn1a* upregulation begins and generates multiple changes to excitatory cells and the circuits they form from early in postnatal life. How these discoveries may contribute to epileptogenesis remains unclear, but they show that pyramidal cells must be considered in future discussions of how the epileptic network emerges in DS.

## Materials and methods

### Animal use and ethical approval

All experimental procedures were performed in accordance with the UK Home Office legislation (2013) Animals Scientific Procedures Act (ASPA) 1986. Mice were housed under a 12 hour light/dark cycle and had ad libitum access to food and water. All pups were kept housed with their mother from birth until experimental use. Wild type mice were of the C57BL/6 genetic background. Dravet syndrome (DS) mice, 129S-Scn1atm1Kea/Mmjax (mice which are heterozygous for the Scn1a knockout allele) were kindly gifted by Dr Rajvinder Karda, UCL. DS mice, and their wild type littermates, were obtained by breeding C57BL/6 females with 129S1/SvImJ background males heterozygous for *Scn1a (Scn1a+/-*). Data was gathered and analysed blinded to pup genotype. All data is divided between genotype and into 3 age groups – postnatal days 10-12 (P10-12), P14-16, and P18-20.

### Acute Slice preparation

All electrophysiology experiments were conducted in freshly prepared ex vivo brain slices. Pups were anaesthetized via isoflurane (3-4% V/V) inhalation until loss of consciousness could be confirmed via lack of toe pinch reflex, and subsequently underwent cervical dislocation followed by immediate decapitation. Dissection of the brain was performed in ice-cold artificial cerebrospinal fluid (aCSF) containing (in mM): 125 NaCL, 25 NaHCO3, 2. 5 KCl, 1.25 NaH2PO4, 2.5 Glucose, 2 CaCl2, 1 MgCl2, bubbled with carbogen (95% O2 and 5% CO2) for at least 20 minutes prior to use. Post dissection, the brain was immediately submerged into an ice-cold aCSF-filled microtome chamber (Leica VT1200S). aCSF was bubbled with carbogen throughout the slicing. Typically, 4-6 300μm thick hippocampal slices were obtained. Slices were then placed into a carbogen-bubbled holding chamber, heated to ~33°C via water bath, for 30 minutes. The holding chamber was then left at room temperature (22-24°C) for a minimum of 1 hour prior to experimentation. Prior to transfer from the holding chamber to the recording chamber, a cut was made between the CA3 and CA1 subfields to limit recurrent activity resulting from action potential back propagation during electrical stimulation of Schaffer collateral fibres.

### Electrophysiology recordings and analysis

All recordings were obtained from CA1 pyramidal cells, identified by cell body position and morphology, using borosilicate glass pipettes with an electrode resistance of between 4-5MΩ. Slices were submerged in room temperature (22-24°C) aCSF (see above for recipe) saturated with carbogen throughout experimentation. All current clamp experiments were obtained using a K-gluconate based internal solution containing (in mM): 125 K-gluconate, 25 NaCl, 2.5 CaCl2, 1.25 MgSO4, 2.5 BAPTA, 2 Glucose, 1 HEPES, 3 Mg-ATP, 0.1 Na3GTP. All voltage clamp experiments were obtained using a Cs-gluconate based internal solution containing (in mM): 125 CsOH, 125 D-gluconic acid, 8 NaCl, 10 Na-phosphocreatine, 10 HEPES, 0.2 EGTA, 5 Tetraethylammonium-Cl, 4 Mg-ATP, 0.33 Na3GTP, 5 QX-314Br. Internal solutions had a final osmolarity of 290-295mOsm and a final pH of 7.3-7.4. At least 5 minutes were left between break-in and the start of experimentation to allow time for internal solutions to perfuse the cell interior. Capacitance was neutralized via MultiClamp Commander and whole cell access resistance was assessed and monitored via brief application of a 10mV voltage step. Recordings were discontinued if access resistance exceeded 25MΩ or discarded from analysis if resistance or resting holding current changed by over 20% relative to initial values.

### Evoked and spontaneous synaptic currents

Experiments were performed in standard aCSF modified with 50μM D-AP5 (Tocris) to block NMDA receptor-mediated currents. Currents were evoked using a tungsten concentric bipolar microelectrode, placed in CA3-proximal stratum radiatum to electrically stimulate the Schaffer collateral fibres. Stimulation was delivered as a 100μs pulse and calibrated to 120% of the minimum intensity necessary to elicit an EPSC. Stimuli were delivered as paired pulses at both 50ms and 100ms intervals. EPSCs were isolated by holding cells at −70mV, IPSCs were isolated by holding cells at 0mV. Recordings were discarded if IPSC latency was less than 1ms greater than EPSC latency. Evoked EPSC and IPSC properties were calculated from averaged waveforms. E/I ratio was defined as the ratio between EPSC and IPSC maximal amplitude. Spontaneous currents were analysed by calculating the total area under the curve of spontaneous synaptic events. Recordings first segmented to remove the 2 second period after stimulation to minimize the impact of rebound stimulation-evoked firing. Data was then detrended to remove minor deviations in baseline across the recording period and baseline was set to 0. Total charge transferred in the recording period was then calculated by integrating recording current via the composite trapezoid rule.

### Intrinsic excitability

The intrinsic properties of CA1 pyramidal cells were assessed at baseline and post-4AP application through current clamp step and slope injections. Input-output relationships were assessed via 500ms current step injections (−40 to 200pA), repeated 3 times for each cell. Action potentials were counted via automatic detection in a custom-written Python script, defined as peaks crossing a membrane voltage of 0mV. Rheobase and action potential morphology were assessed via 1000ms depolarizing slope injections, rising from 0 to 200pA and repeated 5 times for each cell. Action potential properties were assessed using the initial action potential evoked and were assessed using a custom-written Python script. Sag potentials were assessed by injection of 500ms hyperpolarizing current injections (0 to −200pA). Voltage responses were assessed in a custom-written Python script, with the sag potentials defined as the maximal initial hyperpolarizing potential reached during the first 250ms relative to steady state membrane voltage reached in the final 25ms of step injection.

### 4-aminopyridine (4AP) incubation

Hippocampal slices were divided in two, cut down the midline. An additional cut was placed between the CA1 and CA3 subfields to prevent recurrent epileptiform activity. One slice half was bathed in aCSF modified with 25μM 4-AP (“4AP”) for 3-5 hours or 6 hours whilst the other half was bathed in standard aCSF. Depolarizing current step and slope injections were applied to evaluate the intrinsic properties of CA1 pyramidal cells after prolonged 4-AP exposure. Data is presented as values for each 4AP exposed cell subtracted from the mean value for the corresponding aCSF slice for each mouse. To confirm that 25μM 4-AP was sufficient to increase neuronal activity, spontaneous firing frequency of CA1 pyramidal neurons from transverse slices (P14-16) that were incubated in aCSF or aCSF+4AP. Slices incubated in aCSF were recorded in aCSF 1-3 hours after incubation, slices incubated in aCSF+4AP were recorded in aCSF+4AP 3-5 hours after incubation. Neurons were recorded in whole-cell current clamp mode from RMP and action potentials counted if they crossed 0 mV. Pipettes had a tip resistance of 2-3 MΩ when filled with a K-gluconate based internal solution (in mM 142 K-gluconate, 4 KCl, 0.5 EGTA, 10 HEPES, 2 MgCl2, 2 Na2ATP, 0.3 Na2GTP, 1 Na2Phosphocreatine, pH = 7.25, 285-295 mOsm). A calculated +14 mV liquid junction potential was left uncorrected. The protocol consisted of sweeps of 2.5 seconds, with a 500 ms, −10 pA hyperpolarising step used to monitor changes in access resistance and membrane resistance. APs that occurred during this hyperpolarising step were not counted.

### Temporal integration of synaptic and dynamic clamp conductances

Due to difficulties in applying classic dual-electrode approaches to assess input summation, a single stimulation electrode was placed in stratum radiatum and an artificial EPSP was generated via dynamic clamp, with an AMPAR-like conductance applied as described previously in Morris et al., 2017. In dynamic clamp, the injected current is made to depend on the driving force of the conductance one is trying to replicate such that:

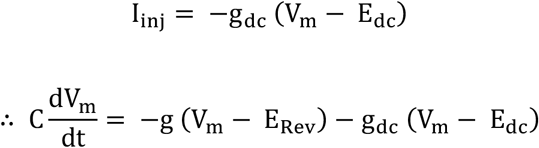

where g_dc_ and E_dc_ are the conductance and reversal potential for the current being replicated. In practice, this must be done via a rapidly implemented loop in which Iinj is adjusted as V_m_ changes. Single channel conductance was set at 0.28nS, reversal potential at 0mV, and a decay constant (τ) of 4ms. Conductance was calibrated by injecting increasingly large AMPAR-like conductances in steps of 0.28nS until firing threshold was reached. This threshold conductance was then reduced by 30%. Stimulation intensity was calibrated by establishing synaptic EPSP latency and introducing the artificial synaptic conductance simultaneous to the start of the EPSP. Stimulation intensity was adjusted until the simultaneous delivery of a synaptic and artificial EPSP had a 50-90% chance of eliciting an action potential (AP). The slice was stimulated every 10 seconds and the artificial EPSP conductance was delivered with an increasing interstimulus interval (ISI), starting at ISI = 0ms and increasing to ISI = 15ms. Throughout recordings, stimulation alone and artificial EPSP alone were delivered every sweep to ensure neither could elicit firing, although a single action potential for each was permitted to allow for the possibility of evoking firing due to summation of spontaneous activity or fluctuations in V_m_. For each cell, 10 sweeps were attempted, but a minimum of 5 was deemed acceptable for analysis. Action potential generation probability was assessed manually.

### *Scn1a* qPCR

Hippocampi from GAD67-GFP mice Tamamaki et al., 2003 at P0, P10 and P20 were extracted from brains and dissociated using Adult Brain Dissociation Kit and OctoMACS Separator (both from Miltenyi Biotec) following manufacturer’s instructions. This procedure collected a large fraction of different cell populations ranging from immune cells to neurons within the same preparation. To purify neurons, the Neuron isolation kit (Miltenyi Biotec) was used according to manufacturer’s instructions. Collected neurons were FACS-sorted by a BD FACS Aria Fusion to isolate GAD67-GFP negative neurons (excitatory neurons) from GAD67-GFP positive (GABAergic interneurons). RNA extraction from GAD67-GFP negative fraction and from total neurons, was performed using TRIzol™ reagent (ThermoFisher Scientific) according to the manufacturer’s instructions. For qRT-PCR, cDNA synthesis was performed using the ImProm-II Reverse Transcription System (Promega). qRT-PCR was performed in triplicate with oligos targeting Scn1a and 18s mRNA using Titan HotTaq EvaGreen qPCR Mix 5x (BIOATLAS). Delta Ct was determined. Primers: 18S_F GGTGAAATTCTTGGACCGGC and 18S_R GACTTTGGTTTCCCGGAAGC; Scn1a_Ex7_F CACCAACGCTTCCCTTGAGG and Scn1a_Ex7_R TGGACATTGGCCTGCATCAG.

### Statistics

All statistical analysis was performed using Prism v6, GraphPad Software, with initial data handling and processing performed using a combination of Microsoft Excel and custom-written Python scripts to automate certain aspects of data processing. For all datasets, significance was defined as a p-value less than 0.05. Unless stated otherwise, all data was assessed by 2-way analysis of variance (ANOVA) followed by Holm-Sidak’s multiple comparisons test. Unless stated otherwise, all datapoints are displayed as mean ± standard error of the mean (SEM).

## Acknowledgements

We thank Dr Jenna C. Carpenter and Irina Zalavina for their help with animal breeding and genotyping, and Dr Rajvinder Karda for the Scn1a +/- mice. We also thank all the researcher in the department for their useful discussions.

