## Supplementary Figures for "Developmental instability of CA1 pyramidal cells in Dravet Syndrome"

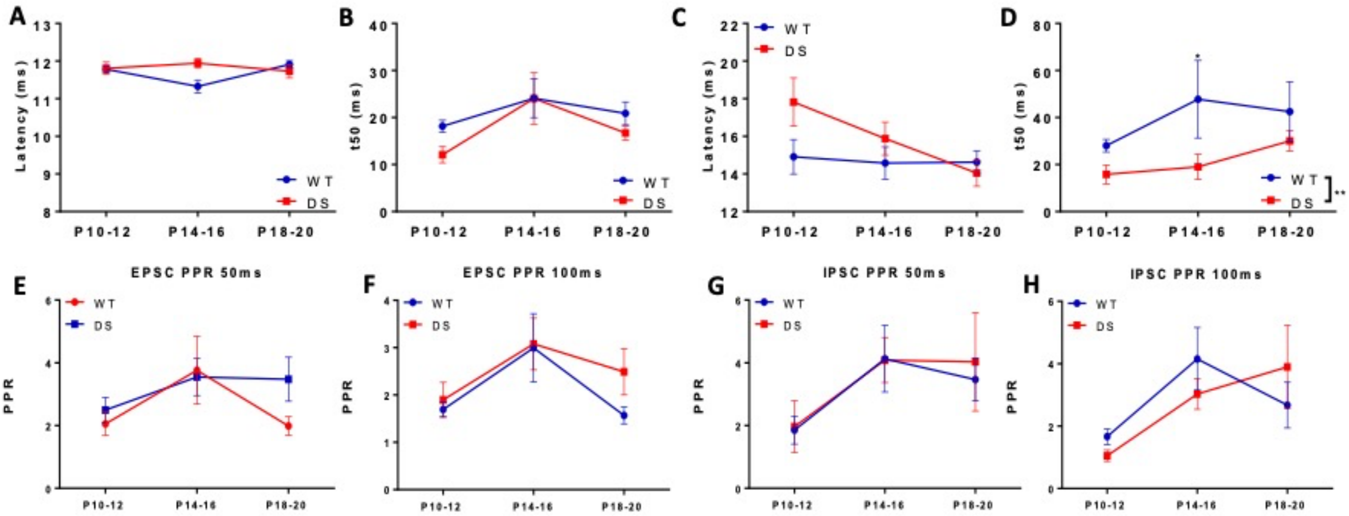


**Figure S1. Secondary parameters of evoked EPSCs and IPSCs.A-D.** EPSC latency (**A**), half-width (**B**), 50ms paired-pulse ratio (**C**), and 100ms paired-pulse ratio (**D**) do not show any inter-genotype differences. Two-Way ANOVA. **E-H.** IPSC latency (**E**), 50ms paired-pulse ratio (**G**), and 100ms paired-pulse ratio (**H**) ratio do not show any inter-genotype differences, whilst IPSC half-width shows a reduction across time in DS mice (**H**, p = 0.008). Two-Way ANOVA. n = WT [31, 16, 12]; DS [10, 21, 30] cells, N = WT [8, 6, 6]; DS [4, 5, 8] mice.

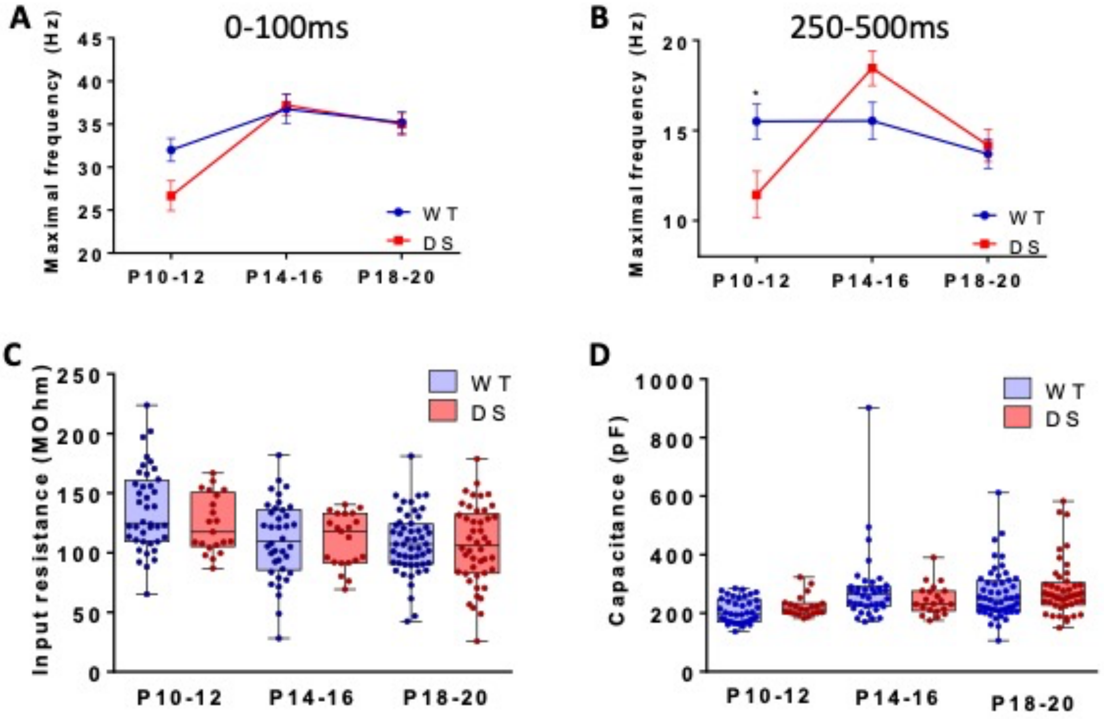


**Figure S2. Maximal action potential frequency, input resistance, and membrane capacitance of CA1 pyramidal cells across DS epileptogenesis. A.** Maximal action potential frequency obtained during the initial 100ms of current step injection show no significant alterations. Two-Way ANOVA. **B.** Maximal action potential frequency obtained during the final 250ms of current step injection show a significant reduction at P10-12 in DS cells. $p = 0.032$, Two-Way ANOVA followed by Holm-Sidak's multiple comparisons test. **C.** Input resistance is unaltered between genotypes across epileptogenesis. Two-Way ANOVA. **D.** Membrane capacitance is unaltered between genotypes across epileptogenesis**.** n = WT [40, 34, 52]; DS [21, 25, 46] cells, N = WT [7, 5, 8]; DS [6, 7, 7] mice.

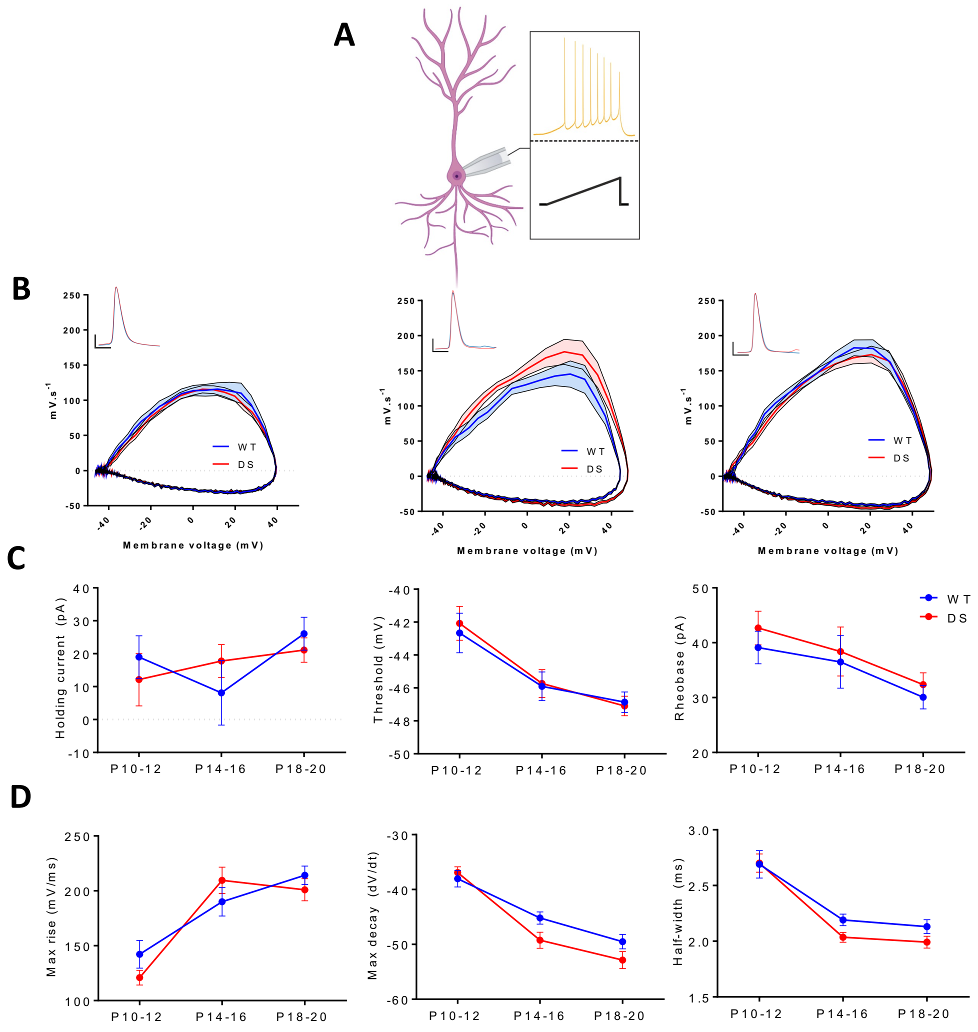


**Figure S3. Rheobase-evoked action potential morphology is unaltered across DS epileptogenesis. A.** Action potentials were evoked at rheobase via current slope injection**. B.** Average action potential morphology and phase plane plots for rheobase evoked action potentials at 3 age groups: P10-12 (left), P14-16 (middle), P18-20 (right). **C.** No significant inter-genotype differences are across epileptogenesis in holding current (left), action potential initiation threshold (middle), rheobase (right). **D.** No significant inter-genotype differences are across epileptogenesis in maximal rise velocity (left), maximal decay velocity (middle), and action potential half-width (right). Two-Way ANOVA. n = WT [34, 28, 48]; DS [21, 29, 39] cells and N= WT [7, 5, 8]; DS [6, 7, 7] mice.

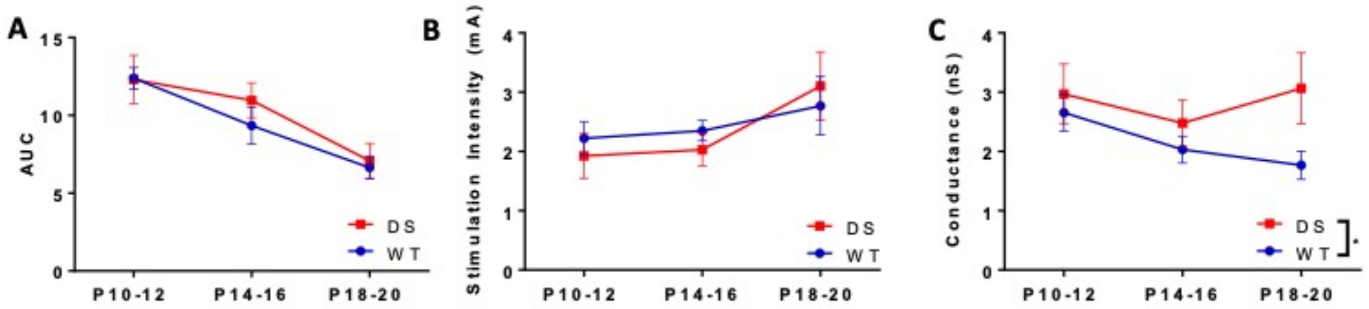


**Figure S4. Input integration is unaltered, but some associated parameters reveal genotype-dependent differences. A.** Area under the curve for integration data action potential probability vs ISI. As with raw data, no genotype effect is seen. Two-Way ANOVA. **B.** Stimulation intensity (mA) used during each integration experiment (**Fig.6**) is unaltered by genotype. Two-Way ANOVA. **C.** Conductance injection amplitude (nS) is significantly altered in DS mice. $p = 0.048$ Two-Way ANOVA. n= WT [11, 8, 19]; DS [10, 16, 11] cells, N = WT [6, 3, 8]; DS [4, 5, 5] mice.

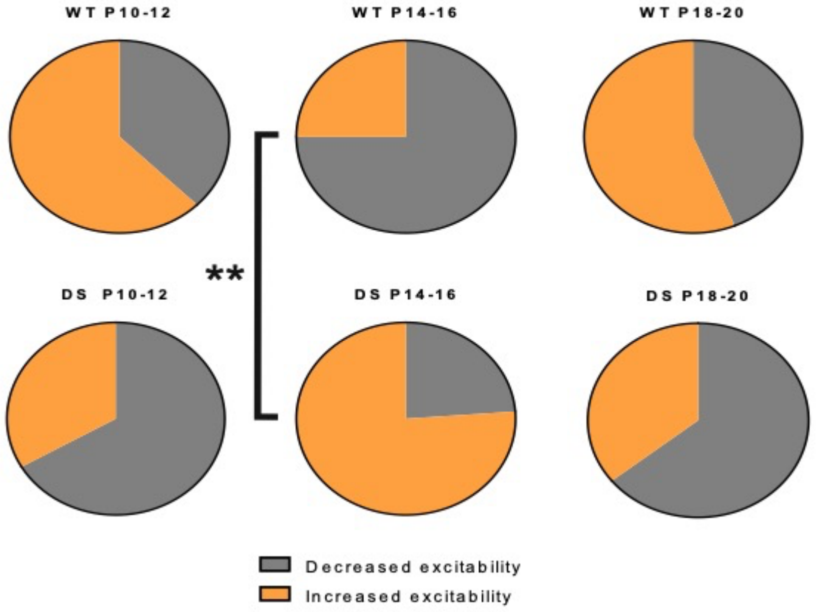


**Figure S5 Proportion of 4AP-exposed cells that increase in excitability relative to control slice excitability.** Proportion of cells that increase *vs* decrease in excitability (maximal firing frequency) in 4AP exposed slices relative to CTRL slice mean. p=0.393, 0.009, and 0.737 for each age group respectively. Fisher's exact test. n = WT [32, 12, 16]; DS [12, 21, 14] cells, N = WT [7, 3, 4]; DS [3, 5, 4] mice.

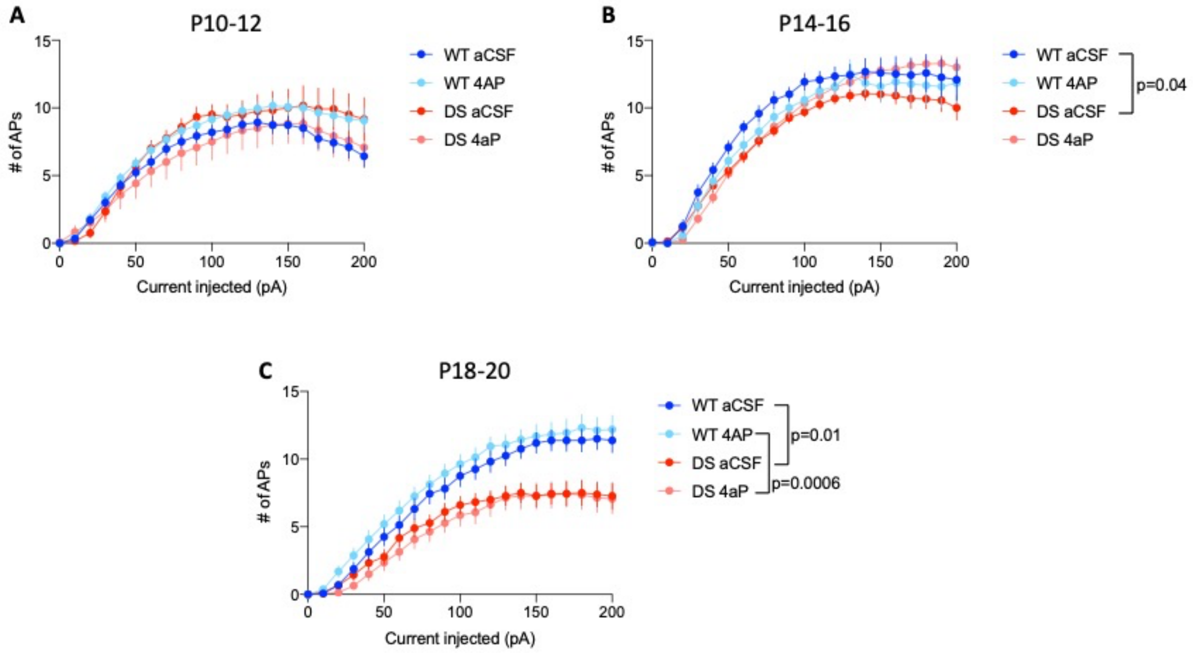


**Figure S6 Raw excitability data from both aCSF and 4AP exposed cells reveals a loss of excitability in DS cells after 6 hours *ex vivo* regardless of treatment group.** I-O curves at P10-12 (**A**) or P14-16 (**B**) and P18-20 (**C**) for WT and DS treated either with aCSF or 4AP. Two-Way ANOVA. n = WT aCSF [30, 12, 16]; WT 4AP [32, 12, 16]; DS aCSF [12, 20, 18]; DS 4AP [12, 21, 14] cells; and N = WT [7, 3, 4] vs [3, 5, 4] mice.

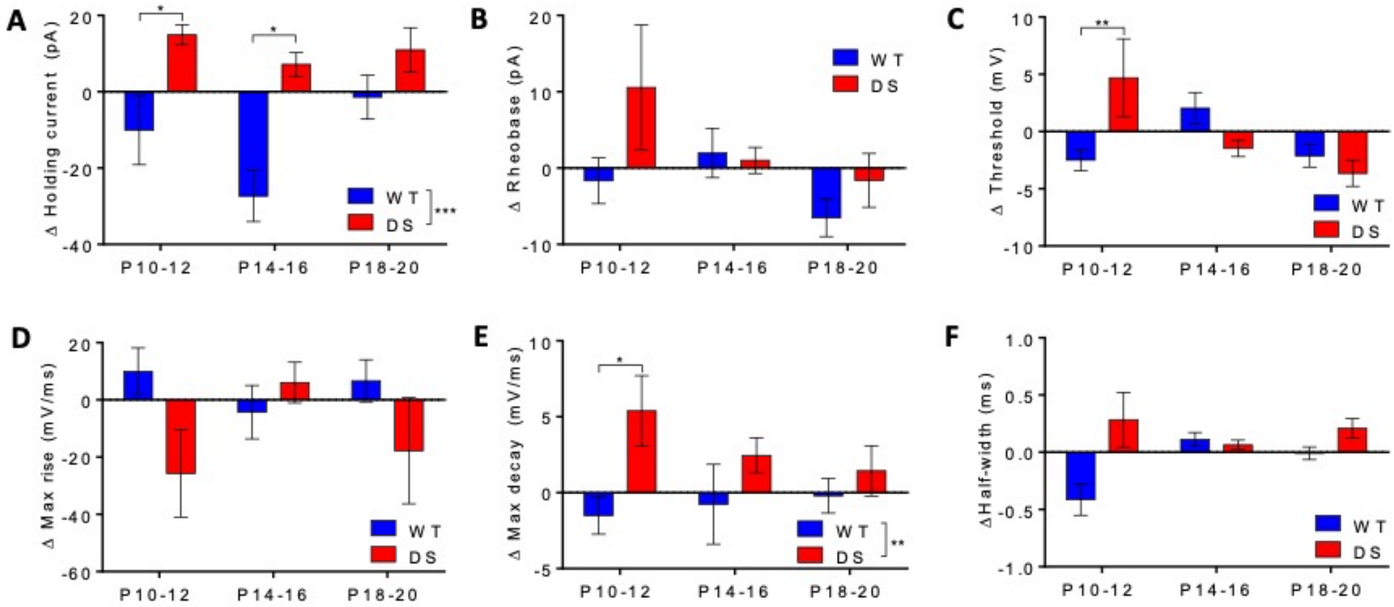


**Figure S7. Intrinsic membrane properties and action potential morphology is differentially altered by 4AP exposure in DS cells. A.** Holding current at $V_m$ = -70mV is consistently increased in DS cells and decreased or unaltered in WT cells post-4AP exposure vs CTRL slices. p = 0.0005, Two-Way ANOVA followed by Holm-Sidak's multiple comparisons test. *p<0.05. **B.** Change in rheobase does not differ between genotypes. **C.** Change in action potential initiation voltage only shows differences at P10-12. Two-Way ANOVA Holm-Sidak's multiple comparisons test. **p<0.01. **D.** Change in maximal action potential rise velocity does not differ between genotypes. **E.** Change in maximal rate of actional potential decay is significantly altered by genotype. p = 0.005, Two-Way ANOVA Holm-Sidak's multiple comparisons test. *p<0.05. **F.** Change in action potential half-width does not differ between genotypes. n = WT [29, 12, 15]; DS [11, 17, 14] cells and N = WT [7, 3, 4] vs [3, 5, 4] mice.
